# Enhancing Biosecurity with Watermarked Protein Design

**DOI:** 10.1101/2024.05.02.591928

**Authors:** Yanshuo Chen, Zhengmian Hu, Yihan Wu, Ruibo Chen, Yongrui Jin, Wei Chen, Heng Huang

## Abstract

The biosecurity issue arises as the capability of deep learning-based protein design has rapidly increased in recent years. To address this problem, we propose a new general framework for adding watermarks to protein sequences designed by various sampling-based deep learning models. Compared to currently proposed protein design regulation procedures, watermarks ensure robust traceability and maintain the privacy of protein sequences. Moreover, using our framework does not decrease the performance or accessibility of the protein design tools.

## 1 Main

Computational protein design, which includes various methods ranging from biophysical models to generative machine learning methods, has emerged as the possible solution to substitute the experimental-based variants selection methods since it could efficiently explore the space of functional protein [1]. Among the computational methods, the capability of generative deep-learning approaches has rapidly grown and can perform various protein design tasks within a single model [2, 3]. Moreover, the experimental validations show that these generative models can design sequences with a high success rate in *in vivo* expression [2, 3]. However, these high success rate protein design tools not only bring opportunities and benefits in biological research but also raise concerns about biosecurity [4, 5], because bad actors in the community could use advanced open-source generative protein design tools to design and synthesize hazard proteins and causes potential threats.

To mitigate this issue, the previous consensus involved controlling the DNA synthesis step and logging every sequence with the authority (*e*.*g*., International Gene Synthesis Consortium, IGSC) for screening to reduce the risk (Fig. 1a) [4]. If *de novo* designed harmful sequences escape from screening due to their little sequence similarity in the database, authority could robustly trace back the source through sequence alignment methods even if a sequence collected from environment may have some mutations. Thus this conventional method can deter bad actors to achieve biosecurity. Although the regulation method is practicable, reporting every sequence to the authority may compromise the privacy of the designed protein sequence and pose challenges to intellectual property (IP) protection. Recently, a concurrent work by SecureDNA foundation introduces a cryptographic framework designed for the screening step to protect the privacy [6]. In this framework, biological sequences are dissected into fragments and then hashed for exact matching with hashed hazardous sequences in a curated database. However, using hash function as encryption method presents difficulties for robust traceability, as the mutated sequences would not have the same hash code as the original one. Additionally, the hashes of all subsequences of length *x* need to be uploaded for screening. These hashes are significantly easier to reverse than a single hash function, which could compromise privacy. As long as a bad actor knows a subsequence *x*_*i*_, · · ·, *x*_*i*+*k*_, they can guess the nearest neighboring base pair *x*_*i*+*k*+1_ with the help of *hash*(*x*_*i*+1_, · · ·, *x*_*i*+*x*+1_). Once the bad actor has the hash of all subsequences and obtains the content of any subsequence, for example, through a collision with a common structure, they can recover the entire sequence with only *O*(*n*) number of hash requests, where *n* is total sequence length. Hence, there still exists a large gap between the current DNA synthesis regulation methods and the need for privacy and biosecurity.

**Fig. 1.**
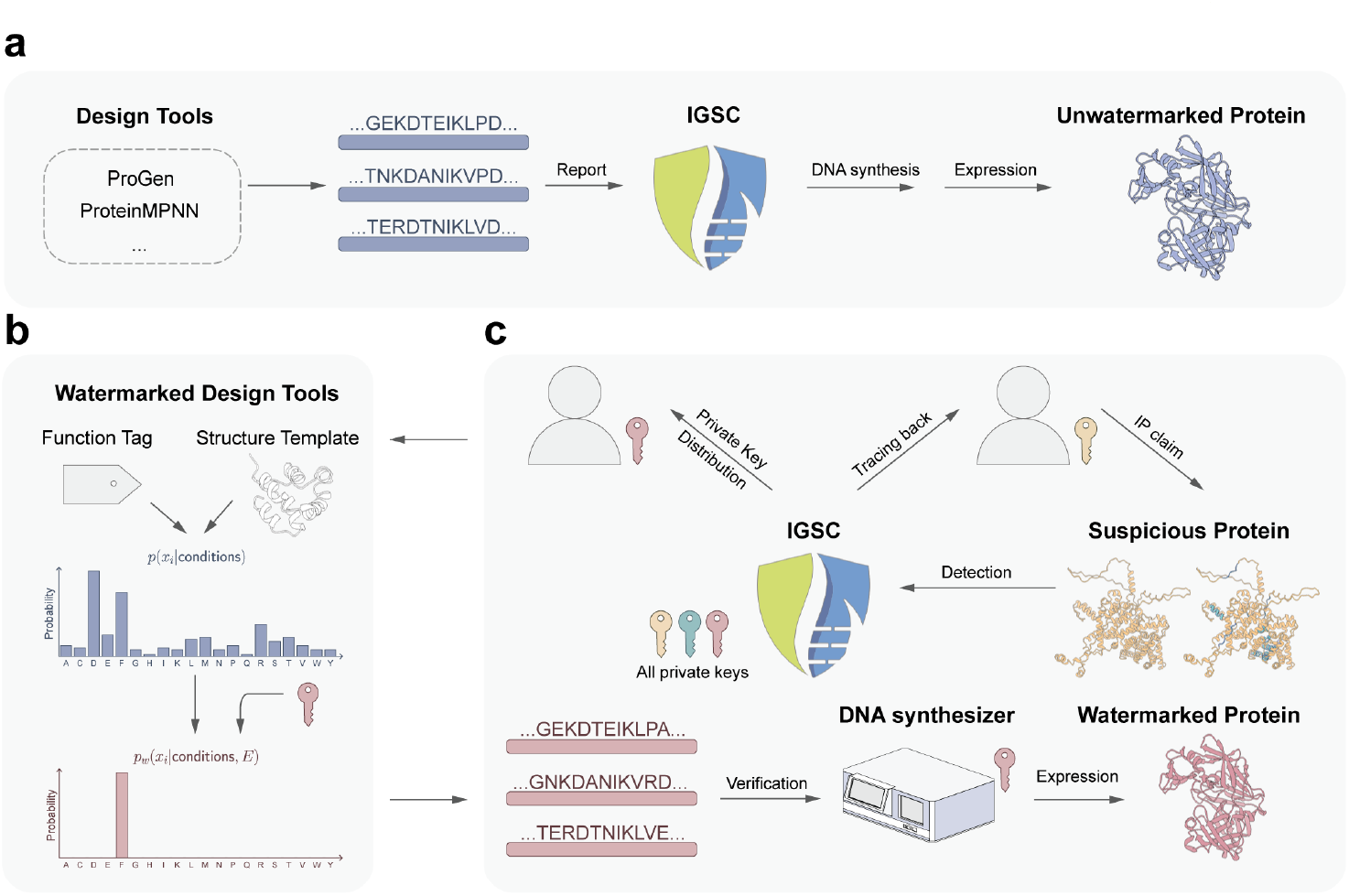
The biosecurity-enhanced regulation process with watermarked protein design. **a)** Conventional protein design regulation process. Researchers use diverse tools to design protein sequences and translate proteins into DNA sequences, then they report every sequence to the authority and obtain permission for synthesis and expression. **b)** Watermarked protein design tools. To add watermarks to protein sequences, the residue distribution of a specific design task is modified according to a watermark code *E* derived from a private key. The expectation of the modified distribution over the watermark code is identical to the original distribution. **c)** Watermarked protein design process. Each researcher applies for a private key from the authority. With this private key, a researcher can design protein sequences with watermarks. These sequences are then translated to DNA sequences and can be locally verified by a DNA synthesizer with a watermark detection program. When a suspicious protein gains interest, the authority could sequence it and trace it back to its designer even it has some mutations. Additionally, researchers could claim the IP of a specific protein based on their private key.

In this paper, we introduce protein watermark, a new general framework for adding watermarks in the generative protein design model to fulfill the need of privacy and robust traceability (Fig.1b,c). In this framework, each researcher gets an authorized private key associated with their identity from the authority, then the researcher can use this private key to add watermarks to the generative model-designed protein sequence (Fig.1b). With the private key and corresponding sequences, the DNA synthesizer equipped with a watermark detection program could locally verify that the sequences are from an authorized user without logging any sequence-related information to the server (Fig.1c). The watermarks establish a strong association between the synthesized sequences and the researcher’s identity. When a suspicious protein is noticed, authority may trace the source by matching the watermarks against all private keys (Fig.1c). The watermarks are robust and not easy to be erased from the design, thereby improving the traceability of each synthesized sequence and deterring the bad actors to enhance biosecurity. Additionally, this framework allows researchers to claim intellectual property rights over synthesized sequences. If the watermarks are not enough in the designed sequence (*i*.*e*. detected watermark score is below a threshold), the researcher could still report the sequence to the authority as an alternative way to get synthesis permission (Fig.1a).

The protein watermark framework is based on previous works that can add watermarks to contents generated by autoregressive models [7–9] and is modified to improve the efficiency of watermark detection. Previous studies have shown that adding watermarks to the content can either be biased, that is the watermarks can be detected by everyone but the model performance will drop, or be unbiased, which means the watermarks are undetectable unless the private key is given while the model performance remains the same. Considering both the performance requirements and the traceability needs of the secure protein design settings, we employ the unbiased watermarking technique in our framework [8]. Specifically, the intuition of the unbiased property can be simplified as: For a given generated distribution *P* (*x*_*i*_|conditions) of the *i*th residue, we modify the distribution to *P*_*w*_(*x*_*i*_|conditions, *E*) based on a watermark code *E* derived from the private key (Fig.1b). Then we ensure the expectation of the modified distribution is equal to the original one: 𝔼_*E*_(*P*_*w*_(*x*_*i*_|conditions, *E*)) = *P* (*x*_*i*_|conditions). That is, the performance would not be changed and the sequence’s watermarks can not be detected without knowing the correct private key to derive the watermark code (see rigorous proof in Watermark embedding). To detect the watermarks in a sequence, we employ and improve the “model-agnostic” detection method. This method takes the pair of private key and sequence as input and outputs the watermark score and p-value. It does not require the logits information from the generative model, making the detection more flexible and practical (see more details in Watermark detection).

To investigate the basic properties of the protein watermark framework, we assessed it on the representative autoregressive model ProteinMPNN [3, 10]. ProteinMPNN is chosen as the representative because the model can use arbitrary decoding order [10, 11] to generate sequence and it has solid experimental verification. To fairly evaluate the performance, we prepared a dataset containing 60 monomer structures with length ranges from 50 to 650 residues released after the training data cutoff date of ProteinMPNN. To show that our framework is unbiased, we generated 50 sequences for each structure under the original and the watermarked ProteinMPNN model at different sampling temperatures (Fig.2a). The generated sequences’ performance is evaluated by the average pLDDT score predicted by ESMfold [12] since pLDDT is used as a surrogate metric to select generated sequences in the original study [3] (Fig.2a). The results show that adding watermarks to the sequences does not change the performance which is aligned with our theoretical analysis (see Watermark embedding). To show the practicality, we then studied the relationship between the average Shannon entropy per residue and the average detected watermark score per residue to show the efficacy of our framework (Fig.2b). The results show that the detected watermark score increases along with the entropy as watermark can be effectively added when the entropy is high. Moreover, the results show that detecting the watermarks with the “model-agnostic” method proposed in the original paper [8] is not satisfying as the negative watermark score appears when the entropy is low (Fig.2b). We then optimized the detection method and found our new method is substantially improved when the entropy is low, thus making the whole framework more practical (Fig.2b, see details in Watermark detection). To show the robustness and correctness of the watermarks and the corresponding detection method, we investigated the effect of sequence modification. The results display a decayed watermark after mutating all alanine to glycine or mutating all glycine to alanine in a sequence (Fig.2c). Of note, though the water-mark score would decrease under this very strong attack, the robustness enables the authority to track the source of a suspicious protein even if it is largely modified after the DNA synthesis step. Besides, our method guarantees that the watermarks can not be detected by a wrong key (Fig.2c). However, these results also indicate there may exist a trade-off between adding watermarks and the performance as the higher sampling temperature would lead to a lower median performance but more watermarks in the sequence (Fig.2a and Fig.2b). To further understand this observation, we next selected the best average pLDDT score of the 50 generated sequences at each temperature for each structure and showed their relative rank (Fig.2c). Selecting the best pLDDT mimics the process of selecting the best sequence for *in vivo* expression [3]. The findings indicate that the best design does not always come from a lower temperature and the performance between temperature from 0.1 to 0.5 is similar. Combining all the observations together, we interpret them from a probabilistic perspective: as the temperature increases, the variance of the sampling increases, and the performance expands. It means that we may still have a chance to sample a good design at higher temperatures but we would spend higher sampling costs.

**Fig. 2.**
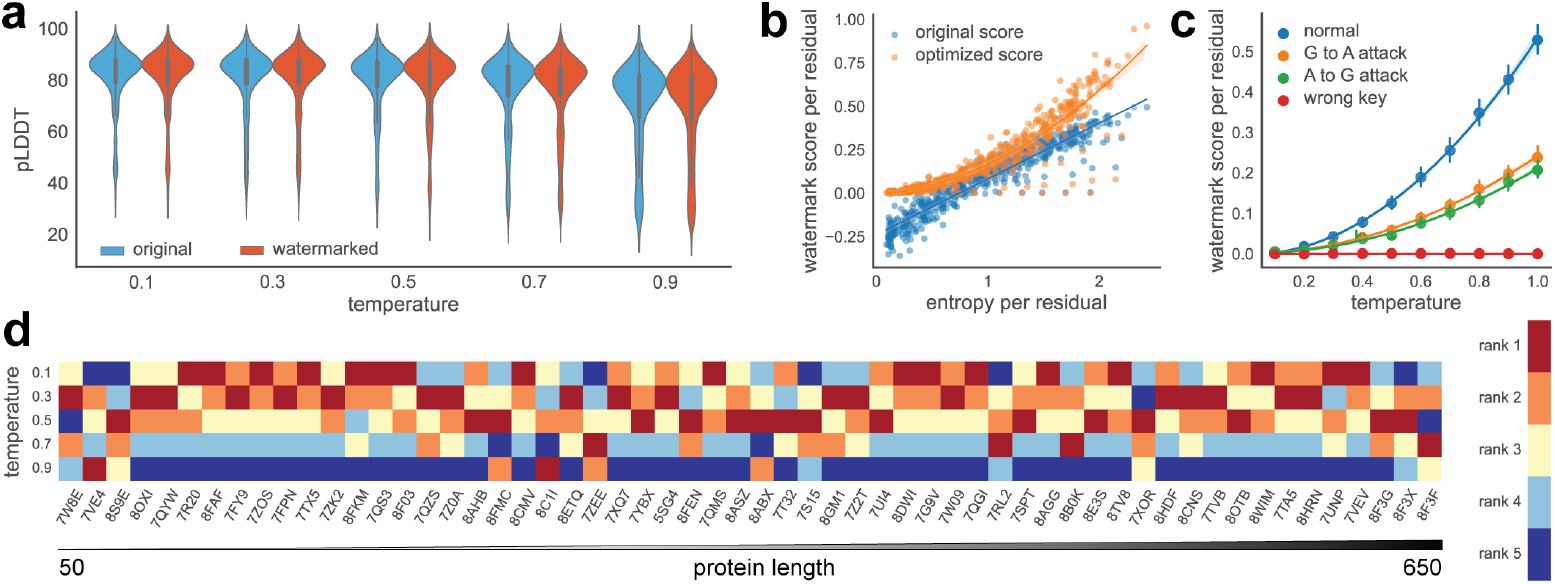
Protein watermark is unbiased, robust, and practical. **a)** The ESMfold’s pLDDT of each designed sequence at different temperatures. The blue and red represent the original ProteinMPNN and watermarked ProteinMPNN, respectively. Each violin plot contains 3,000 pLDDT scores from 60 different monomer design tasks while each task has 50 different designs. The median performance drops as the temperature increases. **b)** The relationship between entropy, and detected watermark score. For the 60 monomer design tasks, we sample one sequence at each temperature. The mean entropy per residue linearly increases as temperature increases. With more entropy, the watermark score detection efficiency increases, and the optimized detection method substantially surpasses the original detection method. **c)** The robustness of watermark score. The average watermark score goes down as the protein sequence is modified. In the plot, “G to A” and “A to G” refer to the substitution of all alanine with glycine and the replacement of all glycine with alanine, respectively. Despite these significant alterations, the watermarks remain detectable. The 95% confidence interval is centered at the mean. **d)** The rank of the best pLDDT score of 50 samples at each temperature. The proteins are ranked based on their length ranging from 50 to 650 residues. These sixty proteins are evenly distributed in terms of length. The designed protein with the highest pLDDT score is in red, whereas the one with the lowest score is in blue.

In summary, the protein watermark is a general, robust, and practicable framework for the regulation of protein design, addressing challenges in privacy need, IP protection, and robust traceability. It can be integrated into the current regulation process and is complementary to hazard sequence detection methods [4, 6, 13]. Based on manipulating the probability space of protein sequences, the protein watermark framework can only be applied to generative models with randomness, and the watermark is hardly detected in low entropy regions. However, these limitations do not conflict with the regulation process, since a low entropy design task means there may be only several proteins in the protein space that can satisfy the requirements, which prevents bad actors from modifying the sequences. Hence, the low-entropy protein design can be supervised by the current regulation process. Furthermore, we envision that more watermarks could be added and detected as the protein functional space could be more efficiently explored in the future. Considering all features together, we believe that the protein watermark framework will play an important role in future protein design regulation processes.

## 2 Methods

### 2.1 Data preparation

In our experiments, we employed 60 protein structures for ProteinMPNN to perform protein redesign task. To make the evaluation reasonable, we searched the monomer structure data with protein length ranges from 50 to 650 released after ProteinMPNN training data cutoff date (from July 22, 2021 to Mar 22, 2024). Then we evenly sampled 60 monomers from all qualified monomers based on its length. That is, for each bin (*e*.*g*., length from 50 to 60, 60 to 70…), we sampled one structure based on the sequence length parsed by the `pdb2fasta.py` script from [Rosetta Commons](https://github.com/RosettaCommons/rosetta/blob/baad48240e60f121d84dc6443bd8745c8d8becdc/source/scripts/python/public/pdb2fasta.py). Of note, in our experiments, we found that the parser behaviour of `pdb2fasta.py` is different from the ProteinMPNN’s parser, and the real output sequence length of ProteinMPNN may differ from the expected length. Specifically, the sequence length of “7WC9” and “8A7Z” parsed by ProteinMPNN exceeds 650 too much and these two proteins’ designed sequences are not evaluated by ESMFold [12] because of the memory constraint.

### 2.2 Protein watermark framework

In this section, we present our methodology for watermarking autoregressive protein design models and detecting the presence of watermarks in generated protein sequences. Our approach allows for the embedding of identifying information into the generation process without compromising the quality of the designed proteins. We also introduce techniques for quantifying the strength of the watermark and efficiently detecting its presence.

#### 2.2.1 Problem formulation

We model the protein design process as an autoregressive model, where each step generates a new token based on the previously designed tokens. In the context of protein sequences, tokens can correspond to the 20 standard amino acids, resulting in a total of 20 symbols in the basic setup. Some encoding methods may utilize additional symbols for each amino acid (see [IUPAC](https://iupac.qmul.ac.uk/AminoAcid/A2021.html)), or multiple amino acid tuples can form a single token based on the byte pair encoding (BPE) algorithm [14]. We abstract the set of possible tokens as a symbol set Σ, and the space of all possible sequences as Σ^*^.

The generation process without watermarking can be modeled as an autoregressive model. Given the previously designed tokens *x*_1_, *x*_2_, …, *x*_*n*_, the probability of the next token is denoted as *P*_*M*_ (*x*_*n*+1_|*x*_1_, *x*_2_, …, *x*_*n*_). The probability of designing a complete sequence is given by:

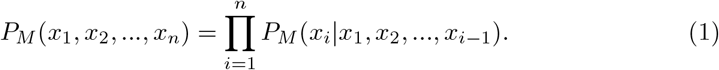

To allow for personalized watermarks, we introduce a key space *R*. Each user selects a key *r* ∈ *R* randomly from a uniform prior *P*_*R*_(*r*). The watermarked generation process is denoted as:

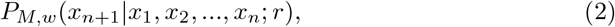

and the probability of generating a watermarked sequence is:

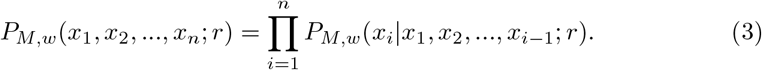

Our goal is to achieve downstream invariance, meaning that for any metric *f* : Σ^*^ → ℝ, the following equality holds:

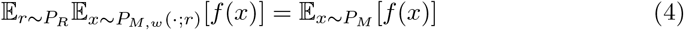

This ensures that the watermarking process does not affect the quality of the generated protein sequences.

#### 2.2.2 Watermark embedding

To embed a watermark, we transform the output distribution in a step-by-step manner. At each step, a new symbol is generated according to some probability distribution over the symbol set Σ. All possible probability distributions over Σ form a simplex, denoted as Δ_Σ_. We introduce a reweighting function *R*_*E*_ : Δ_Σ_ → Δ_Σ_ that depends on a random watermark code *E* following a distribution *P*_*E*_.

##### Unbiased reweighting function

To ensure that the output distribution remains unchanged on average, we require that the distribution is invariant under the reweighting function when averaged over the random watermark code:

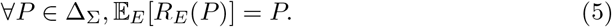

Such a reweighting function is referred to as unbiased reweighting. There exist many instances of unbiased reweighting functions. One example is the *δ*-Gumbel reweighting function, which intuitively uses the Gumbel trick to sample from the original distribution and returns a *δ* distribution over the sampled token as the reweighted result:

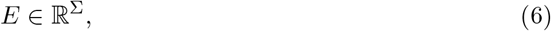

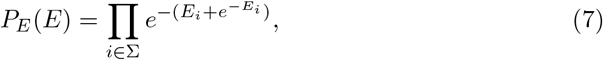

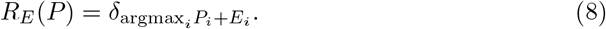

For more examples of unbiased reweighting functions, we refer the reader to [8].

It is important to note that watermarking is only possible for generation processes with sufficient entropy. If the entropy is very low, the reweighted distribution *P*_*M,w*_(·|*x*_1_, *x*_2_, …, *x*_*n*_; *r*) will be very close to the original distribution *P*_*M*_ (·|*x*_1_, *x*_2_, …, *x*_*n*_), making it difficult to embed and subsequently detect the watermark.

In the autoregressive setting, we apply the reweighting function at each step. Given a prefix *x*_1_, *x*_2_, …, *x*_*n*_, the probability distribution over the next token without watermarking is *P*_*M*_ (·|*x*_1_, *x*_2_, …, *x*_*n*_) ∈ Δ_Σ_. At each step, we rely on a random watermark code *E*_*i*+1_, which can depend on the key *r*. With watermark, the new token is sampled from the reweighted distribution:

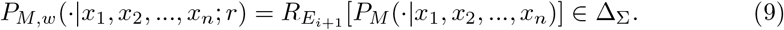

If the reweighting function is unbiased, we have:

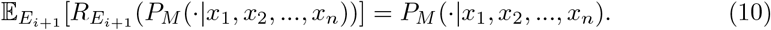

The following theorem shows that if the reweighting function is unbiased and all watermark codes *E*_*i*_ are independent, the entire generation process remains unbiased:

###### Theorem 1.

*If the reweighting function is unbiased and all watermark codes E*_*i*_ *are independent, then* 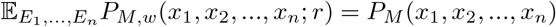.

*Proof*.

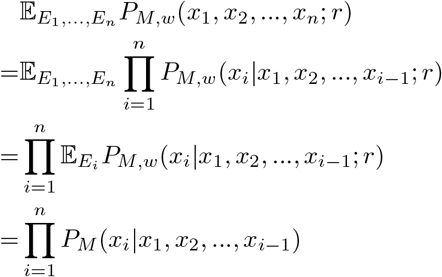

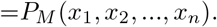

The second equality holds because *E*_*i*_ are mutually independent and *P*_*M,w*_(*x*_*i*_|*x*_1_, *x*_2_, …, *x*_*i*−1_; *r*) only depends on *E*_*i*_. □

This theorem extends the unbiased property of single token generation to the joint distribution of the entire sequence. There exists a natural extension to the generation of multiple sequences: as long as the watermark codes *E* used for generating each token of each sequence are independent, the overall joint distribution of multiple sequences remains unbiased.

##### Generating independent watermark codes

To ensure the independence of the random watermark codes *E*_*i*_, which is crucial for guaranteeing the unbiasedness of the entire sequence generation process, we combine the “context information” and the “key” to construct *E*_*i*_. Specifically, given a prefix *x*_1_, *x*_2_, …, *x*_*n*_, we consider an abstract context code space *C* and construct a context code based on the prefix:

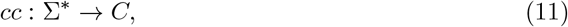

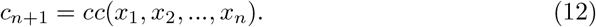

In this work, we use the *m* most recent tokens as the context code:

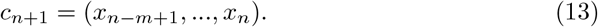

The final watermark code is obtained through a watermark code generation function *Ê* : *C* × *R* → *ε*:

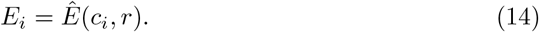

We require the watermark code generation function to satisfy two properties:

- Unbiasedness: In any context, it should generate watermark codes following the correct distribution:

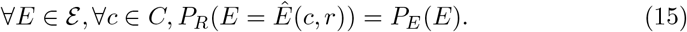
- Independence: For any *n* different *c*_1_, …, *c*_*n*_ ∈ *C*, the generated watermark codes *Ê* (*c*_*i*_, *r*) are mutually independent random variables.

Such a watermark code generation function always exists, as shown in the following proof:

*Proof*. Assume that watermark code space is *ε* and the watermark code distribution is *P*_*E*_(*e*). We construct a key space *R* = *ε*^*C*^, where each key *r* is a function from the context code space to the watermark code space. The random key has a PDF 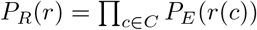. With this construction, *Ê*(*c, r*) = *r*(*c*) constitutes an instance of a watermark code generation function. □

In practice, we simulate the above construction using pseudorandom numbers. Specifically, we use hash(*c, r*) as a random seed to sample *E* from *P*_*E*_ as an implementation of *E* = *Ê* (*c, r*). In this work, we use SHA-256 and a 1024-bit random key *r*.

##### Handling repeated context codes

It is important to note that the context code generation process *cc* : Σ^*^ → *C* is generally not injective. This leads to the possibility of encountering the same watermark code in two steps during the generation of a long sequence or multiple sequences, which violates the independence of *E*_*i*_. To address this issue, we introduce a simple but necessary workaround: if a repeated context code is encountered during the generation of a token, we skip the reweighting step for that token. Specifically, we define a context code history that records all the context codes encountered in the watermarking process using the key *r*. At each watermarking step, we compute the context code *c*_*i*_ as usual. If *c*_*i*_ has not appeared in the context code history, we proceed with the normal watermarking process:

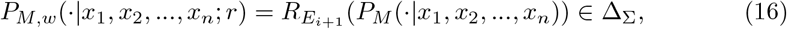

and add *c*_*i*_ to the context code history. Otherwise, if *c*_*i*_ has appeared in the context code history, we skip the reweighting step:

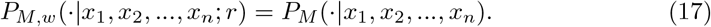

In our experiments, we set the context code length be 5, which means the previous 5 tokens will be used as the context code. Hence, the watermark code space would be 20^5^ = 3, 200, 000 in the experiments. It indicates that the unbiased watermark could be added at most 3,200,000 times for a model, otherwise, the watermark could not be added or could be detectable. In practice, the context code length can be extended to a larger number to satisfy the protein design needs. For example, when the context code length is 10, the watermark space (20^10^) would be large enough to satisfy any protein design needs. Of note, there is a trade-off between context code length and robustness, since a longer context code would lead to a less robust traceability.

#### 2.2.3 Properties of the watermark

The theorem presented earlier has several implications for the properties of the watermark:

##### Preserving output quality

The watermarked and original probability distributions are identical on average:

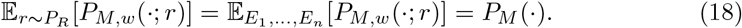

Consequently, the expectations of any metric *f* : Σ^*^ → ℝ are also equal:

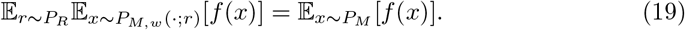

This ensures that the watermarking process achieves downstream invariance and does not affect the quality of the generated protein sequences.

##### Undetectability

For an observer who does not know the watermark code, it is impossible to determine whether a new token is sampled directly from *P*_*M*_ (·|*x*_1_, *x*_2_, …, *x*_*n*_) or from *P*_*M,w*_(·|*x*_1_, *x*_2_, …, *x*_*n*_; *r*) for a random watermark code *E*. It is also infeasible to detect whether a sequence contains a watermark. This property is closely related to the preservation of output quality: if the presence of a watermark could lead to a degradation in quality under certain metric *f*, then there would exist a method to guess the presence of a watermark based on the quality under such metric. Conversely, if the watermark is undetectable, it implies that the output quality is not affected.

However, if the watermark code is known, it is possible to guess whether a token is sampled from *P*_*M*_ (·|*x*_1_, *x*_2_, …, *x*_*n*_) or *P*_*M,w*_(·|*x*_1_, *x*_2_, …, *x*_*n*_; *r*) through statistical tests.

#### 2.2.4 Watermark detection

In the previous section, we discussed the process of watermark embedding, where sampling from the watermarked distribution *P*_*M,w*_(·; *r*) yields a text with a watermark. The detection task is the inverse problem: given a text *x*_1_, *x*_2_, …, *x*_*n*_, we aim to determine whether it is more likely to have been generated from the original distribution *P*_*M*_ (*x*_1_, *x*_2_, …, *x*_*n*_) or the watermarked distribution *P*_*M,w*_(*x*_1_, *x*_2_, …, *x*_*n*_; *r*). This is equivalent to performing a statistical hypothesis test between two hypotheses:

- *H*_0_: *x*_1_, *x*_2_, …, *x*_*n*_ follows the original distribution *P*_*M*_ (*x*_1_, *x*_2_, …, *x*_*n*_).
- *H*_1_: *x*_1_, *x*_2_, …, *x*_*n*_ follows the watermarked distribution *P*_*M,w*_(*x*_1_, *x*_2_, …, *x*_*n*_; *r*).

For each sequence to be tested, we output a real-valued score *S*. By setting a threshold *Ŝ*, we accept hypothesis *H*_0_ if *S < Ŝ*, indicating that there is insufficient evidence to reject the hypothesis that the text was generated by the original model. Otherwise, we reject *H*_0_ and conclude that there is sufficient evidence to support the presence of a watermark.

Under both hypothesis *H*_0_ and *H*_1_, the score *S* and the individual scores *s*_*i*_ are random variables. There are two types of error probabilities associated with this hypothesis test:

- Type I error (false positive): 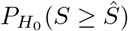.
- Type II error (false negative): 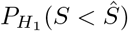.

The p-value of the test is defined as 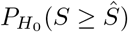, which represents the probability of falsely rejecting *H*_0_ (i.e., concluding that a watermark is present when it is not).

In the following subsections, we discuss two types of scores that can be used for watermark detection while maintaining an upper bound on the Type I error probability.

##### Likelihood-based detection

We first introduce a detection method that is more efficient and can achieve lower p-values, but requires knowledge of not only the sequence to be tested but also the probability distribution of each token in the sequence, i.e., *P*_*M*_ (*x*_*i*_|*x*_1_, *x*_2_, …, *x*_*i*−1_).

Inspired by the Log Likelihood Ratio (LLR) test, which is known to be the most powerful test according to the Neyman-Pearson lemma, we define the individual score for each token as:

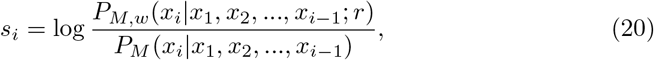

and the overall score as:

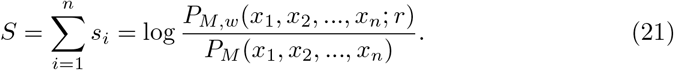

For brevity, we denote *P*_*i*_ = *P*_*M*_ (·|*x*_1_, *x*_2_, …, *x*_*i*−1_) and *Q*_*i*_ = *P*_*M,w*_(·|*x*_1_, *x*_2_, …, *x*_*i*−1_; *r*). Note that *P*_*i*_, *Q*_*i*_, and *S*_*i*_ are all functions from Σ to ℝ, which are equivalent to |Σ|-dimensional vectors. We can define an inner product:

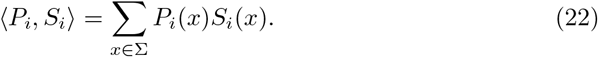

The LLR score can be derived from the following optimization problem:

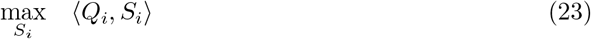

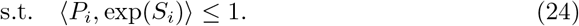

The optimal solution is given by:

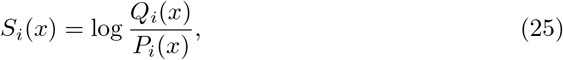

which recovers the optimal log likelihood ratio score *s*_*i*_ = *S*_*i*_(*x*_*i*_).

***Type I error bound***

###### Theorem 2.

*Under hypothesis H*_0_, *we have:*

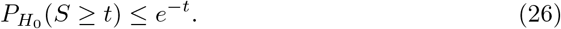

*Proof*. Consider two filtrations:

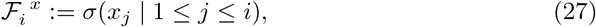

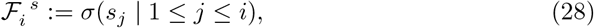

where *σ*(·) represents the *σ*-algebra generated by the random variables.

Due to the constraint ⟨*P*_*i*_, exp(*S*_*i*_)⟩ ≤ 1, the exponential momentum is bounded:

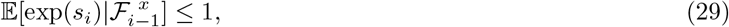

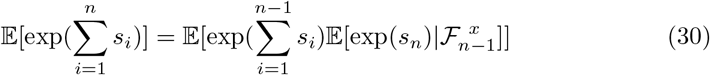

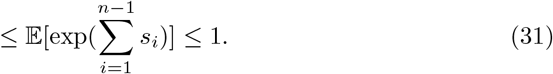

Applying the Chernoff bound yields the desired result. □

Therefore, to ensure that the Type I error probability is bounded by *α*, i.e., 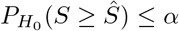, we can set the threshold as:

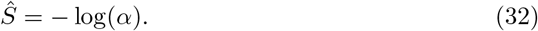

##### Likelihood-agnostic detection

Watermark detection can also be performed in a likelihood-agnostic manner, which does not rely on probability scores for computing the detection score. In [8], a likelihood-agnostic score is introduced for *δ*-Gumbel reweighting function:

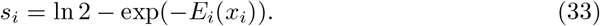

This score also has an exponential momentum bound:

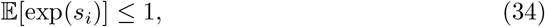

and therefore, it also has a Type I error bound.

However, this score has a large Type II error. In cases where the entropy is not high enough, the score may even be negative, making it impossible to detect the watermark. In practice, this score detects fewer watermarked sequences compared to the likelihood-based score.

To address this issue, we propose an improved likelihood-agnostic score for *d*-Gumbel reweighting function:

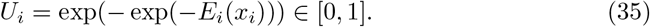

Under hypothesis *H*_0_, *U*_*i*_ follows a uniform distribution on [0, 1]. The log exponential momentum is given by:

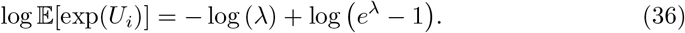

The final score is:

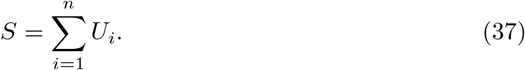

Applying the Chernoff bound, we have:

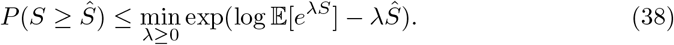

We can obtain the tightest Type I error bound by optimizing over *λ*.

Unlike the likelihood-based detection, the likelihood-agnostic score is not derived using a principled method, and therefore, its design has a certain degree of arbitrariness. The advantage of the likelihood-agnostic score is that it requires less information and does not depend on the probability distribution *P*_*M*_ of the original generation process. This can be more appealing when the original distribution *P*_*M*_ is difficult to estimate.

However, when the original distribution *P*_*M*_ is known, the likelihood-agnostic score tends to be much smaller than the likelihood-based score, leading to worse Type II errors.

### 2.3 Evaluation metrics

To evaluate the quality of designed protein sequences, we followed the previous works [3, 15] and used pLDDT score as the surrogate metric to select promising sequences. We used ESMFold [12] following its tutorial to predict protein structure and the pLDDT score. The reason why we used ESMFold is that ESMFold is very fast and memory efficient compared to AlphaFold2 [16] and it makes the inference of 30,000 sequences affordable in the experiments. Moreover, ESMFold is especially suitable for monomer structure prediction problem and the pLDDT between ESMfold and AlphaFold2 has a good correlation in practice [12].

## 3 Data availability

All data used in this study comes from public database PDB. The PDB structure list is available in our [repository](https://github.com/poseidonchan/ProteinWatermark/tree/bc40bc1c25773694c0c64f7f6276227569b7a711/Experiments/data).

## 4 Code availability

The open source implementation of the protein watermark framework is available at https://github.com/poseidonchan/ProteinWatermark. Detailed tutorials and reproducible experiments are also provided in this repository.

## 5 Acknowledgements

## 6 Author contributions

Y.C., Z.H., and H.H. conceived the idea. Y.C. implemented the main algorithms and conducted experiments. Z.H. proposed and implemented the optimized watermark detection method. Y.C. and Y.J. drew the illustrative figure. Y.W., R.C., and W.C. provided helpful comments on this work. Y.C. and Z.H. wrote the manuscript. All authors revised the manuscript.

